# Temporal Preparation and Statistical Learning: The Influence of Temporal Preparation on Statistical Learning Effects in Complex Temporal Environments

**DOI:** 10.1101/2022.08.06.502990

**Authors:** Josh M. Salet, Wouter Kruijne, Hedderik van Rijn, Eckart Zimmermann, Nadine Schlichting

**Author notes:** corresponding author’s.

## Abstract

Timing studies on statistical learning predominantly present temporal regularities in a discrete, trial-by-trial manner. This, however, is a simplified representation of nature’s complex temporal structure in which regularities are embedded in a continuous stream of interrupting, irregular events. Recent studies using more complex, dynamic experimental tasks have nevertheless replicate the ubiquitous finding of a performance benefit for regular versus irregular timed events. This regularity benefit is commonly interpreted to emerge from a mechanism that detects and subsequently exploits the regularity as it would be impossible for irregular timed events. In contrast, here we show that this interpretation overlooks the influence of learning various time-related factors that contribute to the regularity benefit in such complex temporal environments. Instead of only learning the temporal regularity, participants exploited the temporal properties of targets that we initially considered as irregular and uninformative. Participants temporally prepared, in parallel, for different actions to perform and locations to attend, irrespective of target’s regularity. In fact, this temporal preparation explained regularity benefits without assuming statistical learning of the regular target. Using a computational model, *f* MTP, we illustrate that such adaptation can arise from associative memory processes underlying temporal preparation.

## 1 Introduction

Statistical learning (SL) is a major theme in the cognitive sciences and refers to the learning processes underlying the extraction of statistical regularities in the environment to guide behavior (e.g., Saffran et al., 1996; Sherman et al., 2020). Studies on SL’s temporal aspects investigate whether the environment’s temporal statistics are extracted and used to prepare for when events happen. The ubiquitous finding across a range of paradigms is that, while participants are unaware of any temporal regularity, their performance is better for regular versus irregular timed properties in the experiment (e.g., lower reaction time or higher accuracy in language learning, Hay and Saffran, 2012; Selchenkova et al., 2014; serial reaction time tasks, Olson and Chun, 2001; O’Reilly et al., 2008; Shin and Ivry, 2002; rapid serial visual presentation tasks, Sali et al., 2015; singleton tasks, Xu et al., 2022). These regularity benefits are commonly interpreted to emerge from the ability to extract regular intervals from the environment (e.g., Baker et al., 2004; Boettcher et al., 2022; Fiser & Aslin, 2001; Salet et al., 2021; Sherman et al., 2020; Zhao et al., 2013).

SL’s temporal aspects have particularly been studied in temporal variants of serial reaction time (SRT) tasks (Gobel et al., 2011; Heideman et al., 2016, 2018; Olson & Chun, 2001; O’Reilly et al., 2008; Salidis, 2001; Sanchez et al., 2015; Shin & Ivry, 2002, 2003). In classic SRT tasks (Nissen & Bullemer, 1987), participants make a corresponding keypress to each item in a stimulus sequence (e.g., stimuli A-B-A responded to with index-middle-index finger). In temporal versions of such SRT tasks, a regularity benefit is found when the intervals between successive stimuli contain a regular pattern (e.g., the intervals between A-B-A always follow short-long). That is, although participants were unaware of any regularity, their responses were faster and more accurate to sequences with a regular interval pattern compared to randomly timed sequences. This regularity benefit is only found, however, if the temporal order of the sequences is ordinal (e.g., stimuli are always presented in fixed order A-B-A, O’Reilly et al., 2008; Sanchez et al., 2015; Shin & Ivry, 2002).

Recently, studies have cast these findings with respect to the demands of nature’s complex temporal structure (Boettcher et al., 2022; Salet et al., 2021; Salet, de Jong, & van Rijn, 2022). In contrast to SRT tasks, regular interval patterns in naturalistic environments almost never happen in isolation, but are continuously interrupted by other behaviorally relevant events. In such settings, SL can not operate through a mechanism directly extracting the regular intervals between ordinal events as in SRT tasks. Instead, to extract regular intervals, a mechanism must monitor them over and beyond other intervening events. To test this, recent timing studies developed paradigms in which responses to targets from the regular sequence are continuously interrupted by equally relevant targets from the irregular sequences (Boettcher et al., 2022; Salet et al., 2021; Shalev et al., 2022). Importantly, the overall ordinal sequence in these paradigms, that is, the sequence of any subsequent target regardless of their location, contained no regularities - neither in its temporal order nor in the intervals between targets. Nevertheless, these studies found responses to regular compared to irregular interval sequences to be faster and more accurate. A regularity benefit interpreted to show that classic SL findings, like the SRT tasks described above, can generalize to more complex temporal environments.

In this study, we assess whether this regularity benefit found in complex temporal settings (Salet et al., 2021) results from learning which location to attend or what action to prepare at regular onsets. We set out to answer this question in two preregistered experiments (https://osf.io/3d6h8; https://osf.io/3b78x) in which we coupled the regular sequence to a unique location or action. However, in contrast with our preregistered hypotheses, we found regularity benefits not to result from SL of regular locations or actions, but from a phenomenon known as temporal preparation (see also Addendum to Salet et al., 2021: Salet, Schlichting, et al., 2022). The reoccurring finding in temporal preparation studies is that longer intervals allow more time for preparation, resulting in faster responses (Los et al., 2014; Niemi & Näätänen, 1981; Nobre & Ede, 2018). Similarly, we found performance to improve as more time passed since target’s previous onset. This was observed irrespective of sequence’s regularity and for both location and action, suggesting that participants were capable of monitoring the timing of three sequences in these two dimensions in parallel.

There have been similar reports regarding the involvement of temporal preparation in SL (Heideman et al., 2016, 2018; Salidis, 2001). These studies found that as temporal preparation is not yet fully developed at short intervals, performance benefits more from SL at shorter than longer intervals. Here, we show that such temporal preparation fully accounts for the regularity benefit without assuming SL of the regular target. In contrast with the common SL interpretation that this benefit arises from only learning the temporal properties of the regularity, our findings suggest that the benefit arises from learning multiple time-related factors inherent to the experimental design, both regular and irregular. By using a computational model (*f* MTP, Salet, Kruijne, van Rijn, et al., 2022), we illustrate how such complex, adaptive behavior can arise from memory processes underlying temporal preparation.

## 2 Materials and Methods

### 2.1 Preregistration

This study’s planned sample size, included variables, hypotheses, and planned analyses were preregistered on Open Science Framework (OSF) prior to any data being collected (https://osf.io/3b78x).

### 2.2 Participants

59 First-year psychology students of the University of Groningen were recruited and received course credits for their participation. The experiment was approved by the Ethics committee of the Faculty of Behavioral and Social Sciences (PSY-1920-S-0360) and informed consent was given prior to participation. We continued data collection until we collected data from 54 participants meeting the inclusion criteria (see section 2.5). Sample size rationale was based on Monte Carlo simulations for linear mixed models with the ‘SIMR’ package in R (Green & MacLeod, 2016), aiming for a 80% detection power of a regularity benefit (https://osf.io/3d6h8).

### 2.3 Stimuli and Task

Participants took part in an adapted version of the ‘Whac-A-Mole’ task (WAM, Salet et al., 2021), implemented in OpenSesame (Mathôt et al., 2012) using the PsychoPy backend (Peirce et al., 2019). Participants were seated in a dimly lit room at approximately 60 cm viewing distance from a 22” LCD monitor (100 Hz, Iiyama MA203DT) and aimed to score as many points as possible by reacting to sudden-onset targets with the correct button press. A video of the task is available at: https://osf.io/2ca8h/.

We introduced a color-response mapping in the WAM task (Figure 1). Three target positions were marked by unfilled circles (r = 1.5 cm). At target onset, the circles were filled for 1000 ms with one of three colors, spaced equidistantly (120*^◦^*) in CIELab (L*a*b*: orange [54.0, 56.4, 47.2]; green [54.0, -31.4, 17.1]; purple [54.0, -43.1, -31.4]). Each color coded for a unique action: a button press with the index, middle, or ring finger. Targets in WAM were presented in a continuous sequence that was not divided into discrete trials, and multiple targets could be presented on screen at the same time (coincide by maximally 500 ms). The key manipulation was that, unbeknownst to the participants, one regular target consistently appeared every 3000 ms, while the other two irregular targets appeared at pseudorandom intervals. Each of the three targets was presented equally often (twenty times). The irregular sequence was designed such that the interval between two subsequent presentations was minimally 1250 ms (250 ms between offset and new onset), and that there was always at least one irregular target between two regular target presentations. Importantly, the sequences were designed such that there was no ordinal structure in the overall sequence of any subsequent target (see Appendix A for more details on the stimulus sequence).

**Figure 1.**
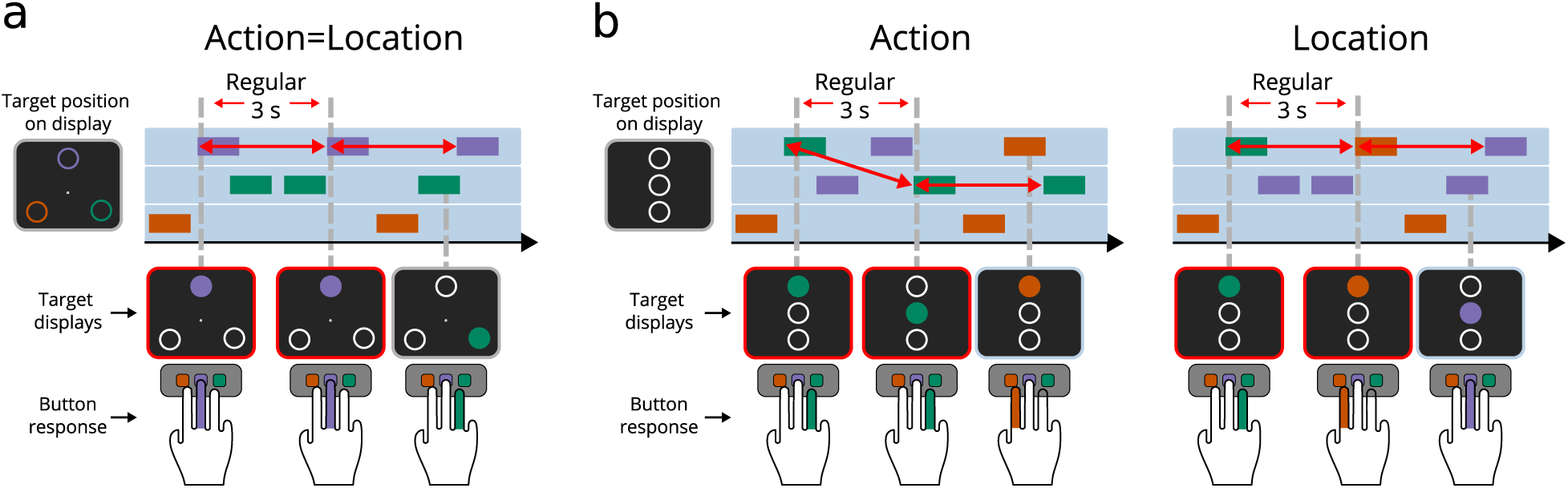
*Experimental conditions.* *Note.* Segments of a series of target presentations in WAM. (a) In the ‘action=location’ condition, action and location were directly coupled and every 3000 ms a regular action was performed in a fixed location (red arrows). (b) In the ‘action’ and ‘location’ conditions, the direct mapping between location and action was removed. In the ‘action’ condition, the regular action was performed every 3000 ms in a random location. In contrast, in the ‘location’ condition, the regular location appeared every 3000 ms and was linked with a random action. The red arrows represent the regular interval, the other events appeared at pseudorandom time points. A video of the task is available at: https://osf.io/2ca8h/.

To assess to what extent SL results from learning regular locations and/or actions, we designed three conditions that differed to what extent the regularity was linked to location or action. In the ‘action=location’ condition, the targets were placed at the three corners of an equilateral triangle (sides = 24 cm). Each target color appeared in a fixed location, requiring a fixed action. One of these was the regular target appearing every 3000 ms (Figure 1a, red arrows). In the ‘action’ and ‘location’ conditions, the direct mapping between location and action was removed (Figure 1b). To avoid Simon interference, the target positions were aligned vertically at the horizontal center of the screen. In the ‘action’ condition, the regular targets required a unique finger response (i.e., all regular targets had the same color), but appeared in a random location (Figure 1b, red arrows). Conversely, in the ‘location’ condition, the regular targets appeared in a fixed location, but had a random color and were thus linked with random actions.

In all conditions, participants scored points by pressing the corresponding color-button on a conventional keyboard with their right index, middle and ring finger (colored stickers were placed on the left, right and down arrow keys). To indicate which button participants pressed, a rectangle in the associated color appeared around the three targets. Points were scored if the correct color-button was pressed within 1000 ms after target’s onset. Each hit was rewarded with three points and accompanied by the sound of a dropping coin. If participants were too late (250 ms after target offset) or pressed an erroneous button, three points were deducted as a penalty.

### 2.4 Procedure

Each of the three conditions consisted of eight experimental blocks in which sixty targets were presented in total, twenty times at each of the three locations and in each of the three colors. A block lasted approximately one minute, and was followed by feedback on the total score, the number of hits and misses, and the high-score. Participants always began the experiment with the ‘action=location’ condition, preceded by one practice block with no regularity. The regular color and location in the ‘action=location’ condition was counterbalanced across participants.

After the ‘action=location’ condition, participants performed four practice blocks with vertically aligned targets where target locations and colors were no longer coupled, without any regularities. The order of the following ‘action’ and ‘location’ conditions was counterbalanced across participants. The features of the regular targets in these two following conditions changed with respect to the ‘action=location’ condition (e.g., if the regular targets were green in ‘action=location’ condition, they changed to orange or purple in the ‘action’ condition). We added the constraint that the middle finger response was never tied to the regularity in the ‘action’ condition, and that the regularity was never in the middle position in the ‘location’ condition. In this way, we aimed to prevent positive biases (i.e., faster and more accurate responses) that we found for the middle finger response and middle target (Supplementary information). At the end of the experiment, participants answered seven questions to assess their awareness of the temporal regularity (reported in Appendix B).

### 2.5 Inclusion Criteria

We excluded and replaced the data from three participants that indicated to be aware of one of the regularities in the questionnaire. We additionally excluded data from blocks in which the hit rate for one of the three targets was less than 25%, suggesting that a participant might not have divided their attention across all targets. Two more participants were excluded because more than three out of eight blocks in one condition had to be discarded.

Response outliers were further identified in two steps: First, a global cutoff (across-participant) was set to remove extreme outliers (only including 150 ms < RT < 1500 ms). Second, we removed RTs that exceed the median -/+ three times the median absolute deviation (computed after the global cutoff) for each participant (within-participant cutoff). All RTs not identified as outliers, were defined as a hit.

### 2.6 Data Analyses

Analyses were conducted in Python 3.8.5 and R 3.6.1 and assessed participants’ responses expressed by RT and HR. We analyzed log-transformed RT using linear mixed models (LMMs, Baayen et al., 2008; Bates et al., 2015) and HR by general (G)LMMs with a logistic link function.

For both RT and HR, we included ‘participant’ as a random intercept. The main interest of the study was whether regularity of targets (regular versus irregular) and their possible interaction with condition (‘action=location’, ‘action’, versus ‘location’) affected performance. As introduced later in section 3.2, we extended the statistical model with the predictor foreperiod (FP). These predictors are reported in the Results section. To investigate order effects, we additionally assessed whether these predictors interacted with the order of condition presentation (‘action *→* location’ versus ‘location *→* action’; note that condition ‘action=location’ was always presented first). Furthermore, we considered the effect of previous response (hit versus miss) and practice effects (time-on-task, indexed by block number). These additional analyses did not reveal any effects on RT or HR (Supplementary information).

We tested the contribution of each of the predictors by means of model comparisons, based on Bayes factors (BF) estimated from BIC scores (Wagenmakers, 2007). To quantify statistical evidence for or against a predictor term, we compare the best statistical model that includes this predictor against the best model without it. For readability, we relegated the details of statistical model selection to the Supplementary information.

## 3 Results

### 3.1 Regularity Benefit

A questionnaire indicated that only three participants were aware of the regularity. Those were replaced by new, unaware participants. The findings of the ‘action=location’ condition (Figure 2a) replicate our previous results of the regularity benefit (Salet et al., 2021): RT was lower and HR higher for the regular compared to irregular targets (Δ*BIC* > 15.7, *BF* > 1000). In our earlier work, we attributed this benefit to a SL mechanism distilling the regular interval from the environment (Salet et al., 2021). For the ‘action’ and ‘location’ conditions (Figure 2b), the HR results align with these findings (Δ*BIC* > 28.0, *BF* > 1000). However, we found no indication of faster responses to regular targets in the ‘location’ condition (Δ*BIC* = 8.9, *BF* = 83.6) and inconclusive evidence indicating slower responses to the regular target in the ‘action’ condition (Δ*BIC* = 1.1, *BF* = 1.7). This inconsistency between HR and RT is surprising: As faster responses lead to higher HRs, we had expected RT to be the strongest driver of regularity benefits (cf. Salet et al., 2021). In the following, we will show that a different explanation of the regularity benefit, rooted in temporal preparation, resolves this apparent inconsistency.

**Figure 2.**
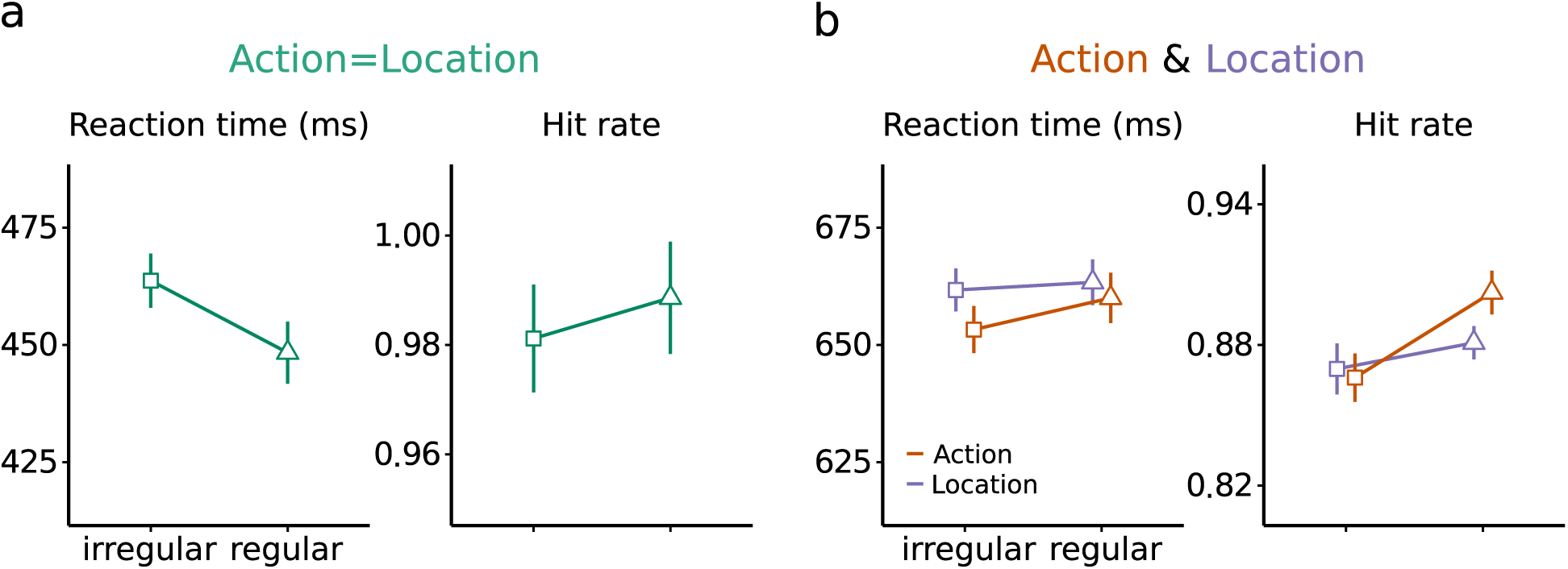
Performance as a function of regularity. *Note*. Mean RT and HR split on regularity with 95% within-subject confidence intervals (Cousineau, 2005; Morey, 2008) for the (a) ‘action=location’ condition and (b) ‘action’ and ‘location’ conditions. Note that the y-scale of ‘action=location’ condition differs from the ‘action’ and ‘location’ conditions.

### 3.2 Temporal Preparation

^1^As the interval between irregular targets was highly variable (1250 – 14000 ms), we have thus far considered it unlikely that these intervals are monitored by participants and used to time responses. However, it has been suggested that participants still monitor these intervals to prepare for upcoming irregular onsets (see Salet, Schlichting, et al., 2022).

#### 3.2.1 Preparing Irregular Targets

In this section, we test whether participants prepare for irregular targets, expressed by a decrease in RT and increase in HR as a function of the interval between each subsequent irregular onset (temporal preparation’s signature: Los et al., 2014; Niemi & Näätänen, 1981; Nobre & Ede, 2018). From here on, we refer to this interval as the foreperiod (FP). Figure 3a illustrates the FP in the ‘action=location’ condition in which action was in correspondence with location. Note that the FP for the regular target was always 3000 ms by definition (Figure 3a, red arrows). In the ‘action’ and ‘location’ conditions, we define the FP as the time between two targets with the same action (e.g., ring-to-ring finger) or location (e.g., top-to-top location), respectively (Figure 3b and c).

**Figure 3.**
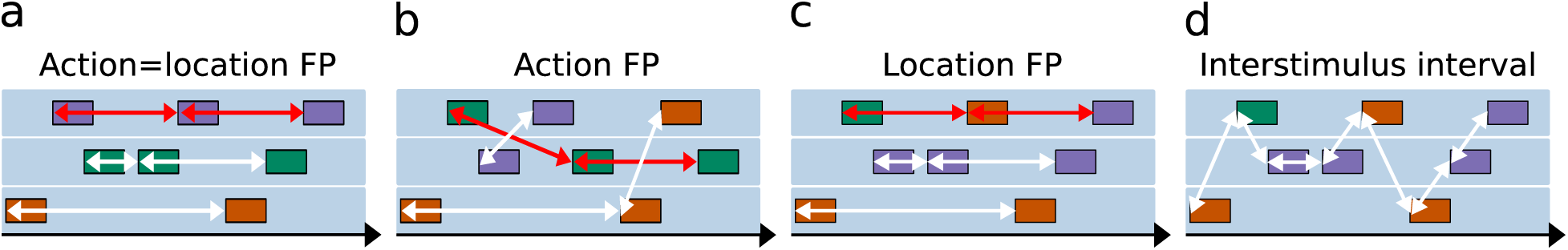
A multitude of temporal properties in WAM. *Note*. Illustrating the foreperiod (FP) in the (a) ‘action=location’, (b) ‘action’, and (c) ‘location’ condition. The red arrows represent the timing of the regular target (constant FP of 3000 ms), the white arrows the irregular targets (FPs ranging from 1250 - 14000 ms). (d) The time between any subsequent target presentation is referred to as the interstimulus interval (ranging from 500 - 2500 ms).

Figure 3d illustrates another temporal property; the interstimulus interval (ISI). This is the time between any subsequent target presentation, and is defined on a much smaller range (500 – 2500 ms) than FPs (1250 - 14000 ms). As can be expected, we found RT to decrease and HR to increase as a function of ISI (Supplementary information). We found, however, that ISI does not modulate the effect of FP on regularity. To focus our report, we relegated these ISI analyses to the Supplementary information and continue with the critical predictors FP and regularity.

To test whether FP duration modulated performance, we extended the statistical model of the section ‘Regularity Benefit’ (predicting RT and HR from ‘regularity’ and ‘condition’) to also include an effect of FP (as a continuous predictor) and its interaction with ‘condition’. Of note, we transformed FP into 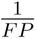 to better capture its curvilinear relation with RT and HR (see Figure 4). This vastly improved the models of both HR and RT compared to a linear relation (Δ*BIC* > 1000, *BF* > 1000). Figure 4 displays RT and HR as a function of binned FP together with model predictions^2^. Model comparisons revealed decreasing RT and HR as a function of FP, suggestive of temporal preparation (Δ*BIC* > 482.7, *BF* > 1000).

**Figure 4.**
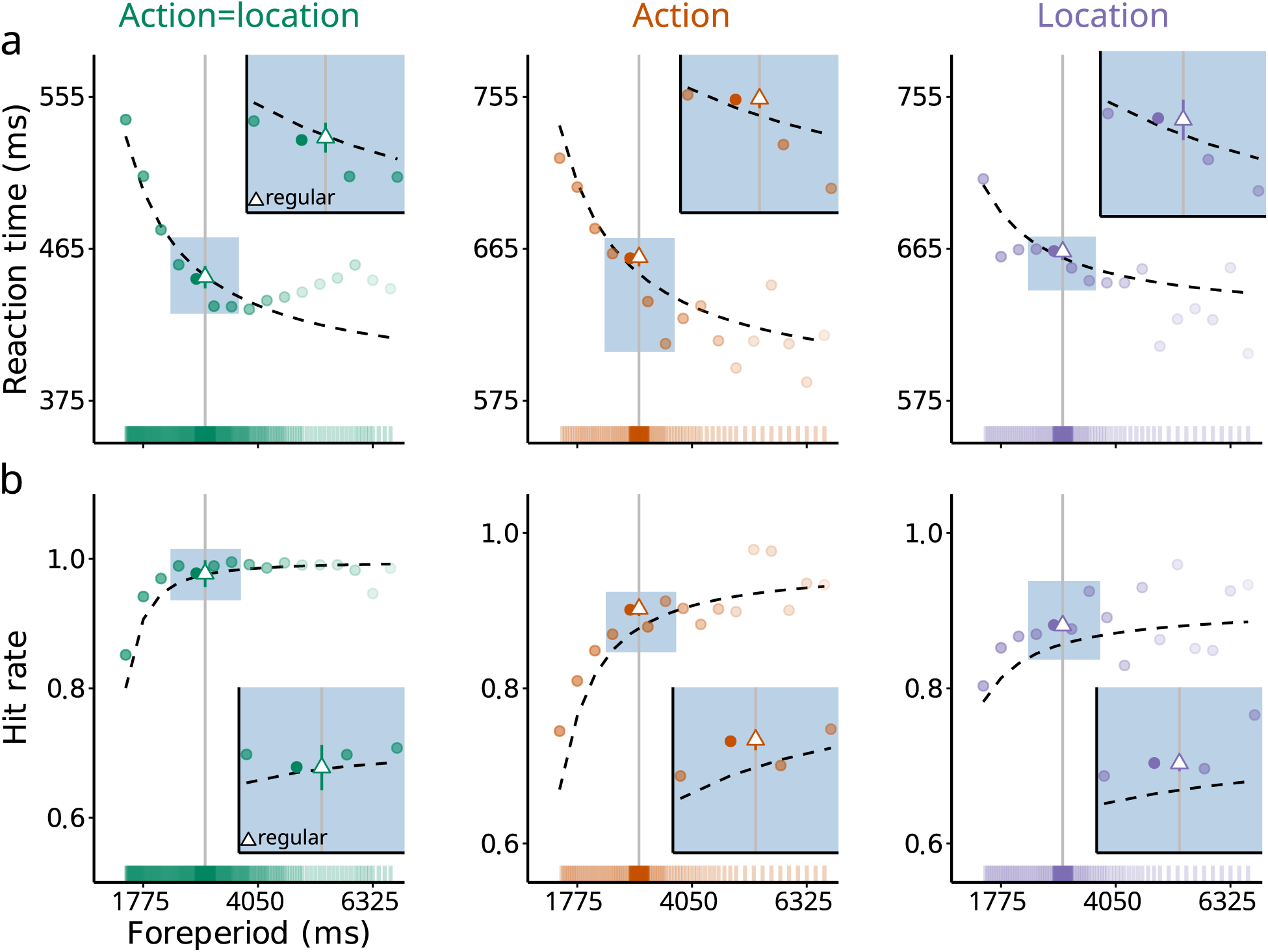
Temporal preparation. *Note*. (a) Mean RT decreases and (b) HR increases as a function of foreperiod (FP), characteristic of temporal preparation effects for both regular and irregular targets. The points represent binned RT and HR (bin size = 350 ms), dashed lines represent statistical model fits. For illustration purposes, we omitted 1% of the tail of the data (FP > 7200 ms). Opacity of the colored bar above the x-axis and the data points illustrate the amount of data per FP bin. Most data is around an FP of 3000 ms (solid vertical line). The open triangle is the same average for regular targets as plotted in Figure 2. As displayed, there is no apparent difference between regular (triangle) and irregular targets (circles) around FP = 3000 ms. Note that for RT the y-axis scale is the same for the ‘action’ and ‘location’ conditions, but differs from the ‘action=location’ condition.

#### 3.2.2 Is the Regularity Benefit an Illusion?

The inclusion of FP as predictor meant that the inclusion of the predictor ‘regularity’ was no longer warranted as fixed effect for both HR and RT (Δ*BIC* > 7.3, *BF* > 38.3). Additionally, the model only including FP vastly outperformed the model including only regularity (Δ*BIC* > 488.8, *BF* >1000). In other words, what seemed to reflect SL can be better accounted for by temporal preparation. This is illustrated in Figure 4, which shows the average RT and HR as a function of regularity and FP. For irregular targets, short FPs (< 3000 ms) are characterized by high RTs and low HRs. Due to the strong asymptotic nature of preparation effects, long FPs (> 3000 ms) are not affected to the same extent. This asymmetry leads to an apparent regularity benefit when only considering the average RT and HR as a function of regularity (as in Figure 2). However, when observing performance as a function of FP, it can be seen that at FP = 3000 ms (Figure 4, insets), there is no apparent difference between regular and irregular targets.

#### 3.2.3 Discussion

We found that participants temporally prepare for all upcoming targets, irrespective of their assigned regularity. Importantly, these findings go against our previous interpretation that participants only capitalize on the regularity embedded in WAM, giving rise to the regularity benefit (see also Addendum to Salet et al., 2021: Salet, Schlichting, et al., 2022). Instead, participants show to monitor both regular and irregular intervals to temporally prepare what action to perform and which location to attend.

### 3.3 What Action and Which Location Comes Next

Thus far, we have defined the FP in the ‘action’ condition as the time between repeating actions, and in the ‘location’ condition as the time between repeating locations (Figure 3b and 3c). This allowed us to evaluate whether FP effects could explain the regularity benefit observed in these conditions. However, each target in the ‘action’ and ‘location’ conditions has both an ‘action’ and a ‘location’ FP (FP_A_ and FP_L_; Figure 5a). It might be that participants exploit both FP types and prepare both what action (e.g., ring finger) and which location comes next (e.g., top location). If so, we should find preparation curves for both FP types in each condition. Additionally, as regularity does not affect performance and the conditions only differed in the assigned regularity (Figure 1b, action versus location), we expect the effect of FP type to be similar across the two conditions.

**Figure 5.**
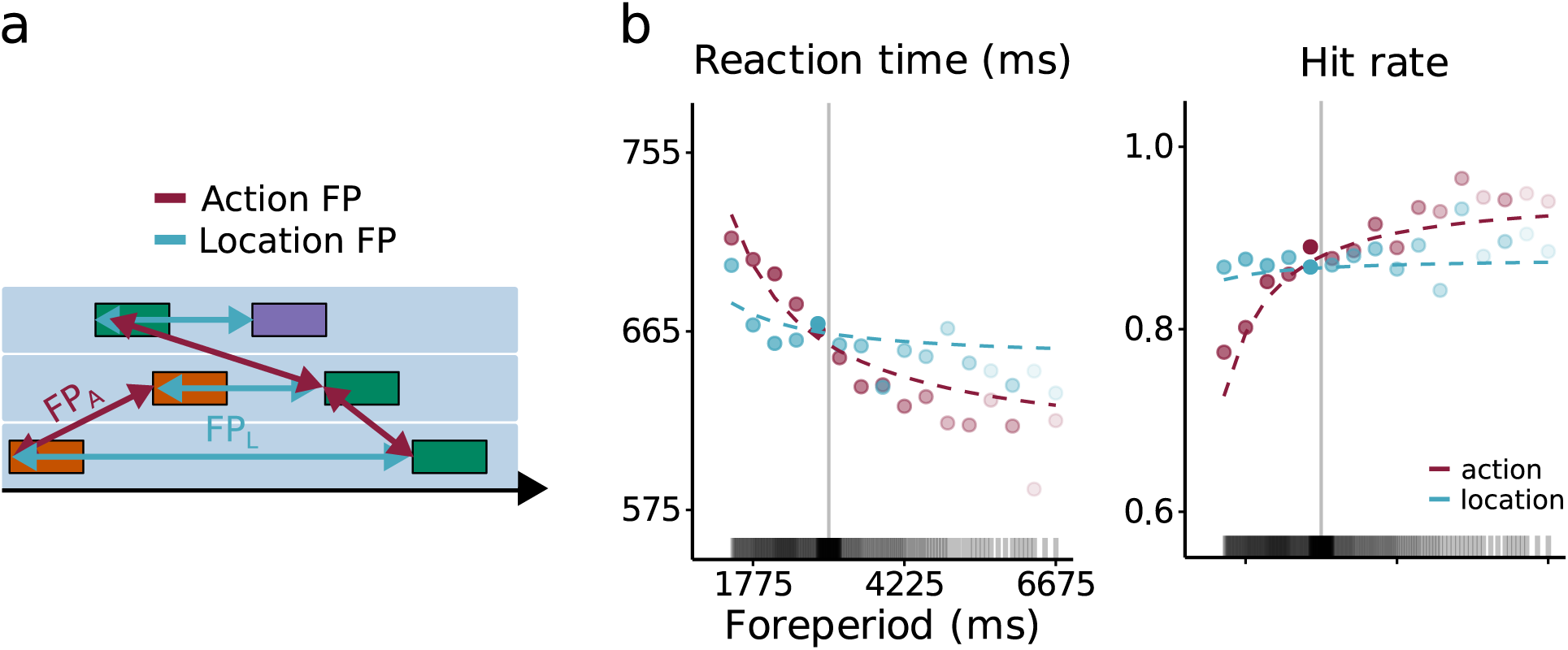
Where and what is next. *Note*. (a) Illustrates the two FP types: action FP and location FP in the same stimulus segment. (b) Mean RT and HR as a function of FP (plotted as in Figure 4). ‘location’ preparation (FPL). In the Supplementary information, we present results of a very similar experiment with 54 participants that replicate all findings presented so far.

To test this, we constructed new statistical models including the predictors FP_A_ and FP_L_, and tested whether they interacted with ‘condition’. Figure 5b displays preparation effects on RT, split on FP type, together with the statistical model’s predictions. Model comparisons revealed an effect of FP_A_ for both RT and HR (Δ*BIC* > 270.3, *BF* > 1000). For FP_L_, an effect was found on RT (Δ*BIC* = 26.0, *BF* > 1000), but evidence of absence for an effect on HR (Δ*BIC* = 6.8, *BF* = 30.3). The models’ steeper RT curves as a function of FP_A_ versus FP_L_ (slope coefficient 4.1 times larger for FP_A_) and the absence of an effect of FP_L_ for HR suggests that preparation effects were more pronounced for action than for location. Finally, FP effects did not differ between the ‘action’ and ‘location’ conditions (Δ*BIC* > 13.9, *BF* > 1000). Indeed, this confirms that the conditions govern similar preparation dynamics and are unaffected by the modality of the regularity (action versus location).

#### 3.3.1 Discussion

Here we show that in the ‘action’ and ‘location’ conditions, participants temporally prepare what action to perform and which location to attend in parallel. Furthermore, we found indications that the effect of ‘action’ preparation (FP_A_) is more pronounced than

### 3.4 *f* MTP: A Computational Account of Preparation

When considering all temporal information embedded in WAM (Figure 3), we find behavioral effects that reflect temporal preparation to both regular as irregular targets (Figure 4 and 5) and not a SL mechanism that only distills the regular interval. In this section, we use a model of temporal preparation (*f* MTP, Salet, Kruijne, van Rijn, et al., 2022) as a proof of concept to illustrate the cognitive mechanisms from which our empirical findings might arise. To do so, we focus on the ‘action=location’ condition and, in particular, on two key aspects of the data. First, does using WAM’s stimulus sequence as input to the model lead to the same preparation curves as observed in the data? Second, does the average RT computed from these preparation effects give rise to an apparent regularity benefit?

#### 3.4.1 Model Architecture

A detailed (mathematical) description of *f* MTP can be found in Salet, Kruijne, van Rijn, et al. (2022). In short, *f* MTP is a neural network consisting of a motor circuit and a timing circuit (Figure 6a). The motor circuit consists of an activation and inhibition node, and their interplay resembles the initiation of a motor plan (i.e., a button press corresponding to the target on screen). During the FP, activation is suppressed by inhibition (Davranche et al., 2007; Duque & Ivry, 2009; Greenhouse et al., 2015; Hasbroucq et al., 1999; Prut & Fetz, 1999; Sinclair & Hammond, 2008), enabling the staging of a motor plan without executing it, akin to an athlete awaiting the start signal before taking off. At target presentation, however, inhibition is released, and the activation node is triggered, initiating the response.

**Figure 6.**
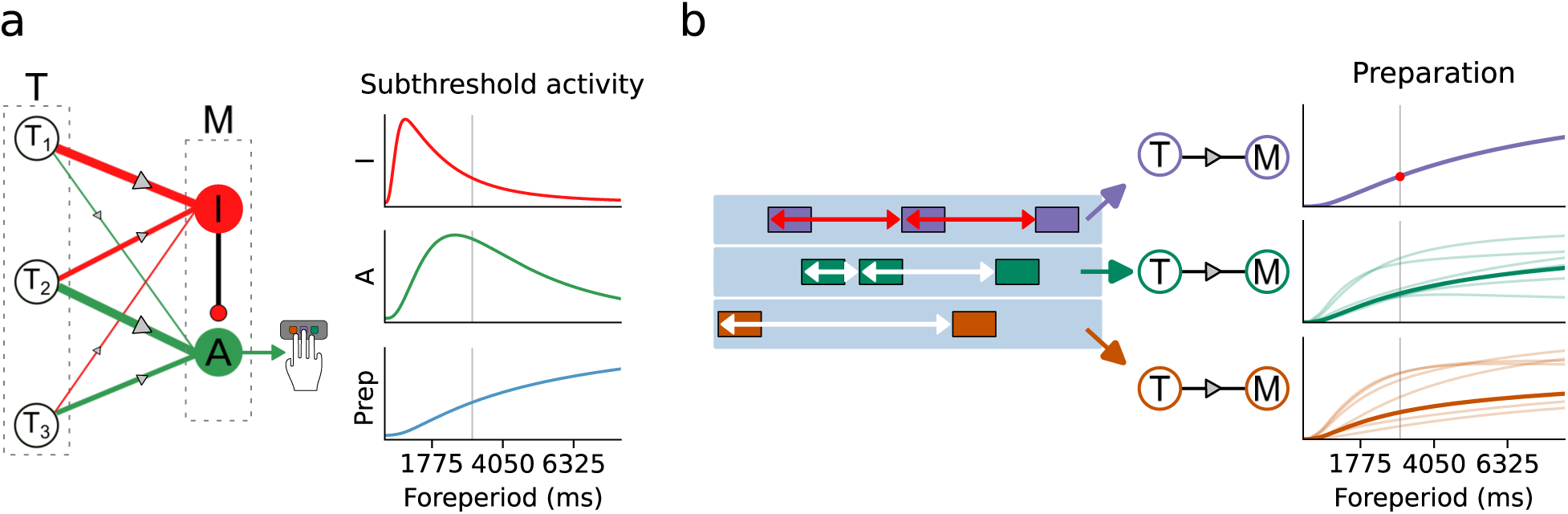
fMTP. *Note*. Illustration of f MTP’s underlying dynamics. (a) Trace formation. In the motor circuit (M), inhibition (I) controls activation (A) to prevent premature responses during the FP. Each node in the timing circuit (T; three example nodes are drawn) projects to both I and A. The thickness of the connections represents the association strength after a sequence of FPs. These associations determine the subthreshold fluctuations in I and A determining preparation (Prep) on the next trial. (b) In WAM, each target (unique location and action) triggers a unique preparation circuit. A constant FP (purple) leads to similar preparation on each trial. As the regular FP is always 3000 ms (solid vertical line), the model’s predicted preparation is only observed at that specific FP (red dot). For the irregular targets (green and orange, varying FPs) preparation fluctuates. Transparent lines represent preparation of different target presentations, the non-transparent line represents average preparation across all presentations. Note that I and A are plotted in arbitrary units, but their ratio (preparation) directly scales to RT.

#### 3.4.2 Trace Formation

This interplay between the inhibition and activation is driven by a timing circuit (Figure 6a, Shankar & Howard, 2012): An array of neuronal units that convey information about the passage of time during the FP (Eichenbaum, 2014; MacDonald et al., 2011; Pastalkova et al., 2008). Different time points during the FP are encoded by different nodes in the timing circuit. Each node in the timing circuit projects to the motor circuit, and their simultaneous firing leads to Hebbian learning (see Gallistel & Gibbon, 2000; Machado, 1997; Mauk & Ruiz, 1992, for similar rules used for learning to time). In this way, specific points in time get associated with either inhibition or activation.

The Hebbian associations between the timing and motor circuit are stored in ‘memory traces’ on each FP trial. A simplified example of this process is illustrated in the left panel of Figure 6a, depicting three example nodes in the timing circuit that encode time points before (T_1_), around (T_2_), and after (T_3_) target onsets. Through Hebbian learning caused by simultaneous firing of the timing and motor circuit, T_1_ primarily forms associations with inhibition (thick red line, it is too soon to respond), T_2_ with activation (thick green line, it is time to respond), and T_3_ neither with inhibition nor activation (thin connection lines, target has already passed). These Hebbian associations define the ‘memory trace’ at each trial.

#### 3.4.3 Memory

Critically, *f* MTP postulates that preparation is determined by the collective contribution of traces formed on previous trials, retrieved from memory. Due to forgetting, traces contribute according to their recency with traces of remote trials contributing less strongly than of recent trials (Wixted, 2004; Wixted & Ebbesen, 1991). It is the contribution of memory traces by which the network’s state tunes to the environment’s temporal properties. The right panel of Figure 6a visualizes the result of this ‘tuning’ after a sequence of multiple FPs (range of 300 - 5500 ms). Early points in time become strongly associated with inhibition, whereas later points in time become more strongly associated with activation. These subthreshold fluctuations in the balance between activation and inhibition during the FP reflect preparation (Figure 6a, blue line). For later time points (FP > 3000 ms), as activation starts to decrease later than inhibition, preparation gradually increases. This gives rise to the asymptotic nature of preparation effects.

### 3.5 *f* MTP and WAM

Originally, *f* MTP was developed and tested as a model of experiments in which FPs (range of 300 - 5500 ms) are presented in isolated trials (Salet, Kruijne, van Rijn, et al., 2022). To model the WAM task with three continuous target streams and a wide range of FPs (1250 – 14000 ms), we extend the model such that each target triggers a unique preparation circuit existing of timing and motor circuits (Figure 6b). With this, we assume specificity of preparation. Each target elicits its associated timing circuit that projects to a corresponding motor circuit, representing a specific response. Aside from this assumption (its rationale is discussed in detail in Appendix C), we keep all other aspects of *f* MTP the same and directly adopt the default parameter values reported in Salet, Kruijne, van Rijn, et al. (2022).

#### 3.5.1 Simulation

To arrive at ‘simulated’ RT, we formalize the interplay of inhibition and activation in *f* MTP (i.e., preparation) as the I/A ratio. High levels of inhibition increase the I/A ratio and slow down responses, and vice versa for high levels of activation. Salet, Kruijne, van Rijn, et al. (2022) showed that this ‘simulated RT’ linearly relates to RT. Here, we focus the simulation on the goodness of the qualitative predictions of *f* MTP (i.e., ‘simulated RT’). Of note, we do not simulate HR but hold the rationale that faster RT

Figure 7 shows that the model’s predictions resemble the data. The RT curve for the irregular targets arises from *f* MTP’s memory principles as earlier time points become, through Hebbian learning, primarily associated with inhibition, whereas for later time points activation is more pronounced. This changing balance, shifting in favor of activation as time passes, yields increasing preparation as a function of FP (see Figure 6), resulting in the downward sloping RT curve. Additionally, when observing simulated RT at FP = 3000 ms, there is no apparent difference between regular (always presented at a FP of 3000 ms) and irregular targets. Due to the asymmetric nature of preparation effects, averaging across regularity leads to an apparent regularity benefit, just as observed in the empirical data.

**Figure 7.**
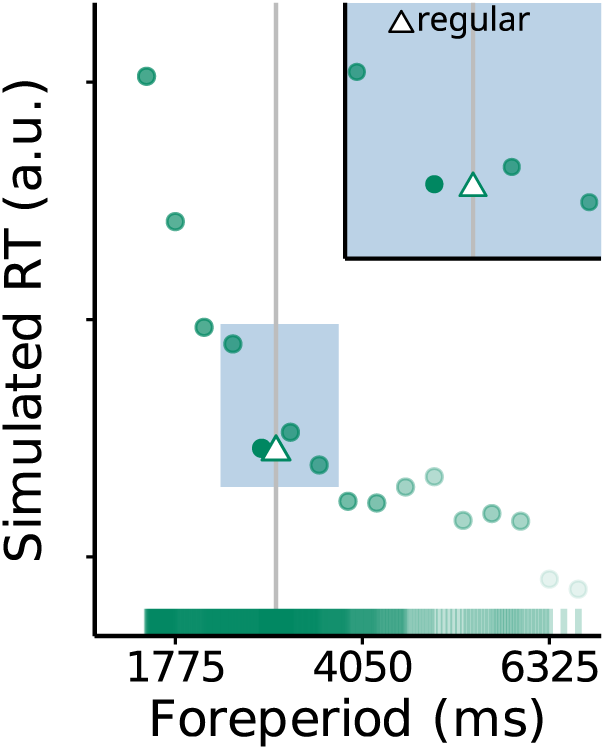
fMTP: simulated RT. *Note*. Mean simulated RT (arbitrary units, a.u.) as a function of FP (plotted as Figure 4 and 5). leads to higher HR, a relation found across all empirical data (Figure 4 and 5).

### 3.6 Discussion

Our simulations show that *f* MTP captures the two key aspects of the data. Guided by *f* MTP, we can make assumptions about the cognitive mechanisms from which our findings might arise. That is, responses in WAM are governed by associative memory processes between timing circuits that monitor the timing of the targets, and motor circuits initiating specific responses. This target-by-target learning process drives preparation during the interval between targets, governing the readiness to respond. In sum, these simulations serve as a proof of concept to show that our results can arise from the memory processes of temporal preparation, without assuming that the processes underlying responses to regular targets are any different from responses to irregular targets.

## 4 General Discussion

SL research is signified by a behavioral benefit for regular versus irregular properties of the environment. This regularity benefit is commonly interpreted as arising from the ability to extract regularities from the environment (e.g., Baker et al., 2004; Boettcher et al., 2022; Fiser & Aslin, 2001; Salet et al., 2021; Sherman et al., 2020; Zhao et al., 2013). In line with this view, we theorized that the regularity benefit observed in WAM’s complex temporal environment is a reflection of a mechanism extracting the temporal information of the regular target (Salet et al., 2021). Intriguingly, however, we found participants to be capable of extracting a range of temporal properties that we initially had considered to be irregular and uninformative. Participants monitored, in parallel, different intervals to temporally prepare what action to prepare in which location, irrespective of target’s regularity. This temporal preparation even showed to fully account for the regularity benefit, leaving the experimental manipulation of regularity in WAM obsolete.

### 4.1 Mechanistic Explanations of Statistical Learning

SL research has convincingly shown the regularity benefit to be a robust finding across different domains (e.g., Fiser & Aslin, 2001; Olson & Chun, 2001; Saffran et al., 1996; Turk-Browne et al., 2009; Yu & Zhao, 2015; Zhao et al., 2013). The theoretical construct of SL, however, has been argued to be underspecified: mechanistic explanations of the cognitive processes that drive SL’s regularity benefit lag behind the robust empirical phenomena (Frost et al., 2019; Thiessen, 2017).

#### 4.1.1 Experimental Challenges of Finding Mechanistic Explanations

An experimental issue for finding mechanistic explanations of SL is that it is difficult to nail down what experimental factor drives the regularity benefit. Embedding a (statistical) regularity in the experiment can introduce other experimental factors that may contribute to the regularity benefit. An example of this issue has been highlighted for ‘probability cueing’ (Jiang, 2018; Shaw & Shaw, 1977), a visual search paradigm where regularity benefits can be observed for targets that appear in regular compared to irregular locations. It has been argued, however, that these benefits are for a large part driven by repetition priming, i.e., a phenomenon that refers to an improvement in performance when stimuli are repeatedly presented (Maljkovic & Nakayama, 1994; Rabbitt et al., 1979). By having targets appearing more regularly at certain locations (i.e., the manipulated statistical regularity), they almost invariably appear more recently at that location as well (i.e., repetition priming). It has been shown that repetition priming can fully account for regularity benefits, without assuming SL of the regular locations (Kabata & Matsumoto, 2012; Maljkovic & Martini, 2005; Walthew & Gilchrist, 2006).

Similarly, we show that temporal regularity benefits can be fully accounted for by temporal preparation, without assuming SL of regular intervals. Such temporal preparation might similarly play a role in recent work assessing the learning of regular interval patterns in a complex visual search task (Boettcher et al., 2022; Shalev et al., 2022). In this search task, regularity benefits were found and interpreted to arise from SL of the regular spatiotemporal pattern. However, the experiment’s complex stimulus sequence embeds a multitude of other temporal properties just as in WAM (similar to Figure 3). Therefore, as in this study, the regularity benefit might be (partly) driven by temporal preparation to these temporal properties.

Temporal preparation and SL are, however, not necessarily mutually exclusive factors that drive regularity benefits. Just as later probability cueing studies showed that both repetition priming as SL contribute to regularity benefits (Goschy et al., 2014; Jones & Kaschak, 2012), it is possible that this is also the case in complex temporal environments like in our WAM task and Boettcher et al. (2022)’s search task. For example, although we found no evidence for SL, it might be that in different settings of WAM (e.g., a speeded WAM version with lower accuracy scores), the higher predictability of the regular compared to the irregular targets is used to improve performance further.

Likewise, our findings do not imply that regularity benefits found in studies using simpler stimulus sequences, like temporal SRT tasks (O’Reilly et al., 2008; Shin & Ivry, 2002), can be fully explained by temporal preparation without assuming SL. They strongly urge, however, to consider the role temporal factors other than the manipulated statistical regularity play in learning behavior. Indeed, this has already been acknowledged in some temporal SRT studies (Heideman et al., 2016, 2018; Salidis, 2001). In these studies, a larger regularity benefit for shorter compared to longer intervals was interpreted to reflect an interaction between temporal preparation and SL: As temporal preparation is not yet fully developed at short intervals, SL has a stronger impact at shorter than longer intervals. All together, future experimental research could further address to what extent and under what circumstances temporal regularity benefits emerge from temporal preparation, SL, or both.

#### 4.1.2 Statistical Learning as General Memory Processes

Despite these difficulties of dissociating SL of the regularity from learning other experimental factors, one line of theoretical work has put forward a unifying explanation of the learning principles underlying SL’s regularity benefits. Various computational models developed under this view showed that regularity benefits can arise from general memory processes underlying both regular and irregular input (French et al., 2011; Kruijne & Meeter, 2015; Mareschal & French, 2017; McClelland & Elman, 1986; Perruchet & Pacton, 2006; Perruchet & Vinter, 1998; Schapiro et al., 2017; Thiessen & Pavlik Jr., 2013). Although these theories have their differences, conceptually they work from two main principles of memory: activation and a forgetting process modeled as decay or interference. In these memory models, each presented item (irrespective of its assigned regularity) is automatically stored in a memory trace that decays over time. Trace activation increases as an item is repeated, while other items may cause interference and decrease trace activation. In this way, repeating items in the environment (i.e., regularities) gain dominance in memory through associative learning between incoming input and memory traces. The application of these models has provided support that such memory processes can lead to adaptation to the environment’s statistical regularities. Our findings, demonstrating that regularity benefits can emerge from memory processes underlying temporal preparation, instantiate this perspective in the temporal domain. Similar to memory models of SL, we showed that *f* MTP’s associative memory processes indirectly give rise to regularity benefits in WAM.

## 5 Conclusions

In conclusion, we found that temporal preparation instead of SL drives regularity benefits for regular intervals in complex temporal environments. Instead of only extracting a single regular interval through SL, we found participants to extract both regular and irregular intervals to temporally prepare for what action to perform and which location to attend. Guided by *f* MTP, we illustrated that such adaptive behavior can emerge from associative memory processes underlying temporal preparation. In a broad sense, we consider our findings as a warning of interpreting regularity benefits to only arise from learning the manipulated statistical regularity in the experiment. Especially in more complex, dynamic experimental tasks striving to reflect some demands of naturalistic settings (such as WAM), it is not unlikely that participants pick up on other experimental factors that contribute to or drive the observed regularity benefit.

## Data Availability

Experiment files, analysis scripts, full dataset, and model scripts can be found on OSF https://osf.io/2ca8h/.

## Appendices A Stimulus Sequences

For each participant and experimental block, a unique sequence of intervals for the irregular targets was generated. Note that the exact same sequences were used between conditions: sequences only differed in their assigned regularity between conditions, their timing properties remained identical.

The generation of the sequences followed a stringent pseudo randomization procedure. The randomization of the timing of the irregular sequences was minimally constrained such that the interval between two subsequent targets in the same location was minimally 1250 ms (250 ms between offset and new onset), there was always at least one irregular target between two regular targets, and that target presentations at different locations could coincide by maximally 500 ms.

Additionally, the interval between the onset of a target and the onset of any next target (ISI, Figure 3d) was controlled for, such that the distributions of ISI’s preceding regular and irregular targets was equated. Systematically longer ISI’s in either condition could drive the regularity benefit effect in the same direction as SL. To minimize ISI differences between regular and irregular targets we assessed the ISI distributions for regular and irregular targets by means of the Kullback–Leibler (KL) divergence (a procedure developed in: Salet et al., 2021). For each participant’s stimulus sequence we computed the KL divergence as a measure of divergence between the probability distribution of regular versus irregular sequences. We only used sequences with a KL-divergence lower than 0.001 (see section ‘Interstimulus Interval’ at https://osf.io/3d6h8 for more details).

## B Questionnaire

At the end of the experiment, we assessed participant’s awareness of the regular interval pattern using the following seven questions: (1) They were encouraged to report anything they might have found ‘noteworthy’ in the experiment; (2) After being informed about the temporal regularity in the ‘action=location’ condition, they were asked whether they had noticed this regularity (Yes/No); (3) They were presented with a forced choice question to indicate (or guess) which target location/color had been the regular one; (4) After having been informed of the existence of the temporal regularities in the ‘action’ and ‘location’ condition, they were presented with questions to indicate whether they had noticed these (Yes/No); (5) They were asked a question to indicate in what order the regularities had been presented; Lastly, they were presented with two more forced choice questions to indicate (or guess) the color (6a) and location (6b) of the temporal regularity.

In the open question, none of the participants reported to be aware of the regularity. In the forced-choice yes/no questions, 5.5% (‘action=location’ condition) and 13.0% (‘action’ and ‘location’ conditions) indicated to have been aware of the regularity. However, only 30.0% (‘action=location’ condition) and 34.0% (‘action’ and ‘location’ conditions) of participants were able to correctly identify the regular color and/or location, closely in line with a guessing rate of 33.3%. Three participants were replaced because they both indicated to have been aware of the regularity, and had correctly identified the regular color and/or location.

## C Specific versus nonspecific preparation

In our modelling (section 3.4), we characterized preparation in WAM as a specific process, meaning that each target presentation triggers its associated timing circuit that projects to a corresponding motor circuit representing a specific response (Figure 6b, main text). This is in contrast with earlier work in which *f* MTP characterized preparation as a general, ‘non-specific’ process independent of different response options (Salet, Kruijne, van Rijn, et al., 2022). A version of the model that incorporates this assumption, however, fails to capture preparation effects observed in WAM (Supplementary information). This mismatch motivated us to adopt the ‘specific’ preparation model presented in the main text. As shown in Figure 7 (main text), this model’s predictions qualitatively align with the data. Although direct empirical evidence for this assumption is as of yet lacking, our results fit within a broad range of behavioral and neurophysiological studies which address the (non)specificity of motor preparation.

Traditionally, the finding that inhibition also applies to task-irrelevant effectors has led to the view that preparation comes with widespread non-specific inhibition of the entire motor system (Hasbroucq et al., 1999; Prut & Fetz, 1999). This is how we had implemented *f* MTP (Salet, Kruijne, van Rijn, et al., 2022), and aligns with the observation that preparation effects are qualitatively highly similar in simple-versus choice RT tasks (Bertelson & Boons, 1960; Frith & Done, 1986; Los & van den Heuvel, 2001; Steinborn & Langner, 2012).

In contrast with this ‘non-specific’ view, however, studies have also found higher inhibition of the selected compared to the non-selected effector of the response, suggesting at least some specificity of inhibition (Davranche et al., 2007; Duque & Ivry, 2009; Greenhouse et al., 2015). This finding has led to the ‘gain modulation hypothesis’ that proposes that inhibition enhances the specificity of selected responses (Duque et al., 2017; Greenhouse et al., 2015). The specificity of inhibition can be illustrated by imagining an ‘inhibition spotlight’ centered around response representations (Greenhouse et al., 2015). A narrow focus, increasing specificity, solely inhibits the selected response, while a broad focus also inhibits non-selected responses. Interestingly, the focus – and thereby specificity – is hypothesized to be context dependent. In tasks with a single response option (e.g., simple RT task), the focus is more broad and therefore also inhibits non-selected responses. However, with multiple response options (e.g., choice RT task), its focus narrows to enhance the separation between the response representations; thereby decreasing the probability of selecting the incorrect response.

In WAM, participants had three response options, and continuously risked a penalty for incorrect choices. From the view of the gain modulation account, it might be that these task demands narrowed the spotlight of inhibition, thereby characterizing preparation as a specific process. A range of behavioral studies have similarly shown that preparation can, at least in some contexts, be specific for the selected response (Langner et al., 2018), target identity (Thomaschke et al., 2011, 2016; Wagener & Hoffmann, 2010), and target location (Wagener & Hoffmann, 2010). These findings align with the specificity assumption made in the main text (Figure 6b). However, to what extent and under which circumstances preparation reflects a (non)specific process remains a question that requires more research.

## Footnotes

1 All following analyses were not preregistered.

2 Note that statistical model predictions represent estimated marginal means (i.e., at the average level of other predictors) and not a least-squares fit to empirical data.

