## Supplementary Information for "Temporal Preparation and Statistical Learning: The Influence of Temporal Preparation on Statistical Learning Effects in Complex Temporal Environments"

### Author Note

Funding: During this project, Josh M. Salet, Wouter Kruijne, and Hedderik van Rijn were supported by the research program "Interval Timing in the Real World: A functional, computational and neuroscience approach", project number 453-16-005, awarded to Hedderik van Rijn, financed by the Netherlands Organisation for Scientific Research (NWO). Nadine Schlichting and Eckart Zimmermann were supported by the European Research Council (project "moreSense", grant agreement 757184).

**Supplementary Information:**

**Temporal Preparation and Statistical Learning: The Influence of Temporal Preparation on Statistical Learning Effects in Complex Temporal Environments**

### S.1 Full Statistical Report

Here, we provide all details of the statistical procedure for the model selection of the (General) Linear Mixed Models (G)LMM to analyze response time (RT) and hit rates (HR). Statistical evidence for or against an effect of interest was quantified by comparing the best statistical model (in terms of BIC) with this effect against the best model without this predictor term. For some comparisons, this means that the compared models can differ in terms of other predictors as well. In the R-notebooks on our OSF repository (https://osf.io/2ca8h/) we list full tables with all possible models ranked by BIC, which determines which models were compared.

#### S.1.1 Middle Bias

As documented in our preregistration (https://osf.io/3b78x), we suspected a 'middle bias' in the 'action' and 'location' conditions. That is, we hypothesized there would be a benefit of (1) detecting targets in the middle location, which was in the center of the screen and (2) a facilitation of middle finger responses compared to the index and ring finger responses. For this reason, these responses were never associated with any regularity by design.

Model comparisons indeed revealed a middle bias: RT was lower and HR higher for middle finger responses ($\Delta BIC = 13.4$, $BF = 801.0$) and for responses towards the middle location (RT: $\Delta BIC = 51.0$, $BF > 1000$; HR: $\Delta BIC = 114.3$, $BF > 1000$). Only for the HR of the middle finger responses, evidence for such a bias was inconclusive ($\Delta BIC = 1.6$, $BF = 2.2$). The best RT models also included a random slope for the effect of 'middle action' and 'middle location' for each participant. For this reason, all analyses in the main text exclude targets presented at the central location or requiring a middle finger response.

We note, however, that analyses including those 'middle' responses yielded the same conclusions.

#### S.1.2 Regularity Benefit

##### *S.1.2.1 Reaction Time*

Model comparisons revealed that there was an interaction between 'condition' and 'regularity' ($\Delta BIC$ = 75.7, $BF > 1000$). This interaction entailed that in the 'action=location' condition RT was lower for regular compared to irregular targets ($\Delta BIC$ = 15.8, $BF > 1000$); in the 'location' condition RT did not differ ($\Delta BIC$ = 8.9, BF = 83.6); and in the 'action' condition we found an inconclusive trend indicating higher RT for regular targets ($\Delta BIC$ = 1.1, BF = 1.7). In addition, there was support for the fixed effect 'condition' ($\Delta BIC$ = 213.6, $BF > 1000$). Figure 2 in the main text, indicate that RT was lower in the 'action=location' blocks. However, we did not perform post-hoc tests to further explore their differences. Finally, there was support for predictor 'previous response' (hit/miss) ($\Delta BIC$ = 142.1, $BF > 1000$). The coefficient of 'previous response' entailed that RTs were higher when the preceding trial ($n - 1$) was missed compared to when it was a hit. Of note, for all following reports of the 'previous response' predictor, the direction of the effect was in the same direction: RT was higher and HR lower when $n - 1$ was a miss. The random effects structure of the model included a random intercept for each participant and random slopes for the effect of 'regularity', 'condition', and 'practice effects' (time-on-task indexed by block number).

##### *S.1.2.2 Hit Rate*

Model comparisons indicated that HR was overall higher for regular than for irregular targets ($\Delta BIC$ = 28.2, $BF > 1000$), but we found no interaction between regularity and condition ($\Delta BIC$ = 7.3, $BF$ = 38.4). Besides support for 'regularity', as reported in the main text, we found support for 'condition' ($\Delta BIC > 1000$, $BF > 1000$) and 'previous response' ($\Delta BIC$ = 449.6, $BF > 1000$). The random effects structure of the best model contained only a random intercept per participant.

#### S.1.3 Temporal Preparation

In addition to the support for the fixed effects reported in our article (‘FP’ and removal of ‘regularity’), there was support for an interaction between ‘condition’ and ‘FP’ (RT: $\Delta BIC = 172.4$, $BF > 1000$; HR: $\Delta BIC = 100.9$, $BF > 1000$) and ‘previous response’ (hit/miss) (RT: $\Delta BIC = 142.1$, $BF > 1000$; HR: $\Delta BIC = 213.6$, $BF > 1000$). The interaction between ‘condition’ and ‘FP’ entails that the preparation curves have different slopes (see Figure 4 main text). However, we did not perform post-hoc tests to explore this interaction further. The random effects structure of the best RT model included, besides an intercept for each participant, a random slope for the interaction between ‘regularity’ and ‘condition’, and the effect of ‘practice effects’ (uncorrelated). The HR model only included an intercept for each participant.

#### S.1.4 What Action and Which Location Comes Next

Besides the support for the fixed effects ‘$FP_A$’, ‘$FP_L$’, (main text) and ‘ISI’ (reported below), there was support for an effect of ‘previous response’ (RT: $\Delta BIC = 27.1$, $BF > 1000$; HR: $\Delta BIC = 393.5$, $BF > 1000$). The random effects structure of the RT model included, besides an intercept for each participant, a random slope for the effect of ‘previous response’ and ‘ISI’ (uncorrelated). The HR model only includes an intercept for each participant.

#### S.1.5 Interstimulus Interval

In this section, we aim to titrate the effect of FP (Figure 3a - 3c), ISI (Figure 3d), and regularity on RT and HR. The analyses in the main text focused on the critical role of FP, leading to the exclusion of the predictor regularity. Here, we consider ISI: the time between any subsequent target presentation (Figure S1a). Similar to FPs, but defined on a much smaller range: 500 – 2500 ms versus 1250 - 14000 ms, ISI modulates performance (Figure S1b): longer ISIs result in speeded responses and thereby increase HR. The focus of these analyses is to consider in depth the possible modulatory role of ISI regarding our conclusions of FP and regularity in the main text.

To focus our analyses, we consider only the predictors regularity, FP, and ISI. Furthermore, we focus on the condition in which we found a reliable regularity benefit: the 'action=location' condition (section 'Regularity Benefit', see main text). To elucidate the role of ISI and FP on the regularity benefit, both separately and simultaneously, we constructed three separate models testing for: ISI, regularity, and their interaction (M1); FP, regularity, and their interaction (M2); ISI, FP, and regularity, including the interaction terms between ISI:FP and ISI:regularity (M3)[1]. We follow the exact same model selection procedure as described in the 'Materials and Methods' section. The R-notebooks on our OSF repository ('ISI in depth analyses', https://osf.io/2ca8h/) provide all details and a step-by-step overview of the statistical analyses.

##### ***S.1.5.1 Results***

The first column in Figure S1b displays RT and HR as a function of binned ISI split on regularity. In contrast to the main text, this figure seems to indicate a regularity benefit across the ISI bins. However, this apparent benefit can be still driven by FP instead of regularity. The FP for the regular targets was always 3000 ms, while the FPs for irregular targets ranged from 1250 to 14000 ms. The apparent regularity benefit may result from collapsing across FP that, due to the asymptotic nature of preparation effects (Figure 4, main text), result in an increase in RT and a decrease in HR for irregular compared to regular responses. Indeed, when selecting a subset of the data with FPs around 3000 ms (Figure S1b, column 2 to 3), correcting the asymmetry in FPs between regular and irregular targets, the apparent regularity benefit seems to vanish.

The statistical analyses support these visual interpretations. For M1, only considering ISI and regularity (cf. Figure S1b, column 1), we found evidence for the inclusion of ISI (RT: $\Delta BIC = 79.8$, $BF > 1000$; HR: $\Delta BIC = 37.0$, $BF > 1000$), but evidence against their interaction (RT: $\Delta BIC = 9.8$, $BF > 137$; HR: $\Delta BIC = 7.8$, $BF =$

[1] Note that the interaction between FP and regularity can not be tested as the regular FP has a single value of 3000 ms.

**Figure S.1**

*Interstimulus interval.*

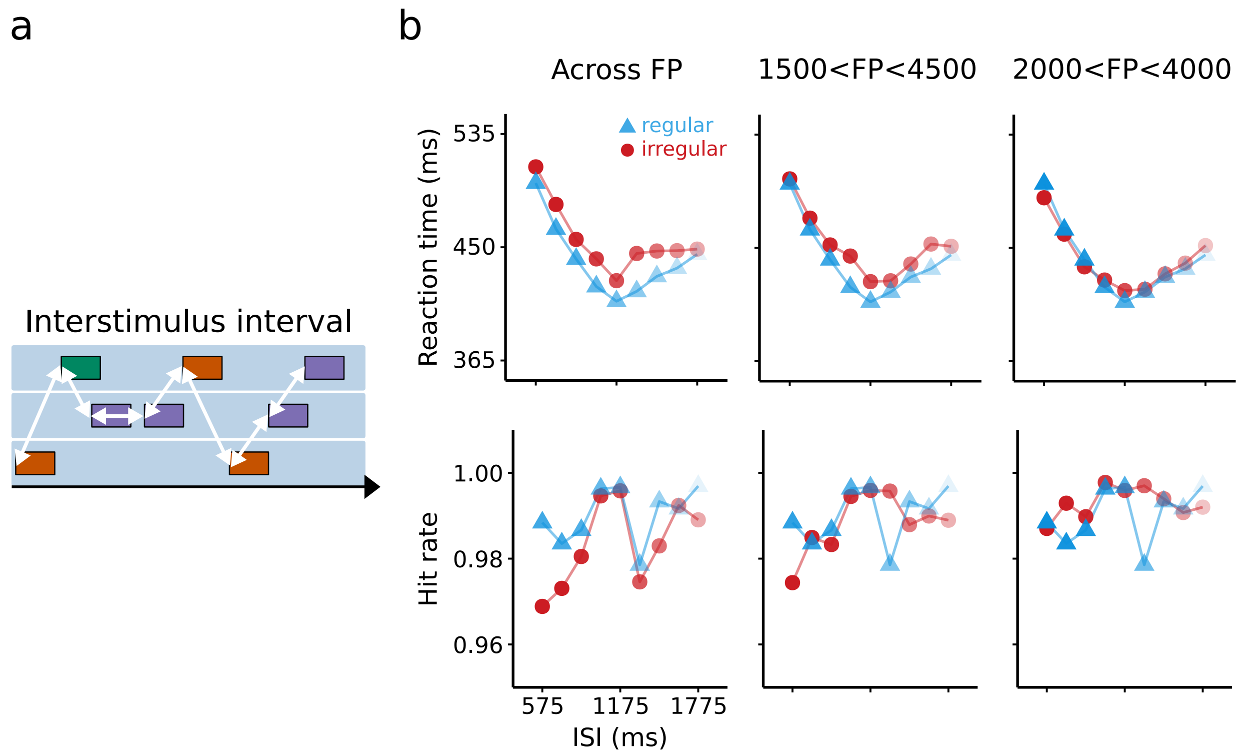


*Note.* (a) Illustrating the interstimulus interval: the time between any subsequent target presentation. (b) Mean reaction time and hit rate as a function of binned ISI (bin size = 150 ms, plotted as Figure 4 and 5). Each column represents different subsets of the data based on the range of FPs: 1250 to 14000 ms (first column, 100 % of data), 1500 to 4500 ms (second column, 80 % of the data), and 2000 to 4000 ms (third column, 58 % of the data). It becomes clear that the apparent regularity benefit when collapsing across all FPs (first column) vanishes when selecting a subset of data with FPs around 3000 ms (second to third column).

48.7). This is indeed in line with Figure S1b (first column) that is suggestive of a regularity benefit when neglecting FP.

For M2, only considering FP and regularity, we found evidence for the inclusion of FP (RT: $\Delta BIC = 135.0$, $BF > 1000$; HR: $\Delta BIC = 311.0$, $BF > 1000$) and evidence against the inclusion of regularity (RT: $\Delta BIC = 5.7$, $BF = 17.0$; HR: $\Delta BIC = 1.9$, $BF = 2.6$).

For M3, considering ISI, regularity, and FP (cf. Figure S1b, column 3), we found evidence for the inclusion of FP (RT: $\Delta BIC = 158.0$, $BF > 1000$; HR: $\Delta BIC = 264.0$, $BF > 1000$), ISI (RT: $\Delta BIC = 90.5$, $BF > 1000$; HR: $\Delta BIC = 10.1$, $BF = 157.0$), and their interaction for RT ($\Delta BIC = 23.6$, $BF > 1000$), but not for HR ($\Delta BIC = 11.6$, $BF = 323.0$). Importantly, we found again evidence against the inclusion of regularity (RT: $\Delta BIC = 2.5$, $BF = 3.5$; HR: $\Delta BIC = 18.6$, $BF > 1000$). The coefficient of the interaction between ISI and FP for RT revealed that the effect of FP on RT grows as a function of ISI. Post hoc Tukey's HSD test revealed that the effect of FP at ISI levels of 500, 1000, 1500, 2000, and 2500 ms were all significant ($p < 0.01$). Evidence for the absence of regularity is in line with Figure S1b (third column), indicating that responses to regular and irregular targets do not differ when selecting the subset of the data around FP = 3000 ms.

##### ***S.1.5.2 Conclusion***

The analyses in this Appendix served as a stringent control analysis of our main findings in the main text. In particular, we aimed to ensure that our conclusions regarding the role of FP on regularity hold when controlling for ISI. Although we found ISI to drive preparatory behavior, it does not modulate the effect of FP on regularity. In sum, we again conclude that FP is the critical predictor that leads to the omission of the effect of regularity. In other words, when we take into account temporal preparation (i.e., predictor FP), we no longer find evidence for a regularity benefit.

### S.2 Replication

The experiment we discussed in the main text is a follow-up of an earlier preregistered experiment (https://osf.io/3d6h8). In this experiment, which is refered to as 'Replication' of the experiment discussed in the main text, we presented participants with 'action' and 'location' conditions. Following Salet et al. (2021), all targets were arranged in an equilateral triangle, and this experiment did not include an 'action=location' condition. In all other respects, the two experiments were identical. As we found no indication of a regularity benefit in either condition of this 'Replication' experiment reported here, and we ran the experiment in the main text as a follow-up, reasoning that the absence of a 'regular benefit' might have been a consequence of Simon interference obscuring any small regularity benefits.

Here, we assess whether the FP effects reported in the main text could similarly be identified in the data of this experiment. As displayed in Figure S2, RT and HR in this experiment were also modulated by FPs. Model comparisons revealed that we replicated all critical effects: We found an effect for both RT and HR of $FP_A$ (RT: $\Delta BIC = 81.6$, $BF > 1000$; HR: $\Delta BIC = 326.9$, $BF > 1000$), $FP_L$ (RT: $\Delta BIC = 130.6$, $BF > 1000$; HR: $\Delta BIC = 7.2$, $BF = 37.4$), and ISI (RT: $\Delta BIC = 60.6$, $BF > 1000$; HR: $\Delta BIC = 22.8$, $BF > 1000$). Additionally, for HR, we found an interaction between $FP_A$ and ISI ($\Delta BIC = 12.5$, $BF = 614.1$). As such, this experiment provides further support of our findings in the section 'Temporal Preparation' of the main text. Finally, we again found support for the fixed effect 'previous response' (hit/miss) ($\Delta BIC = 36.2$, $BF > 1000$; HR: $\Delta BIC = 98.7$, $BF > 1000$). The random effects structure of the model included a random intercept per participant and a random slope for '$FP_A$' (for RT only, not for HR ), 'previous response', and 'ISI'.

**Figure S.2**

*Replication.*

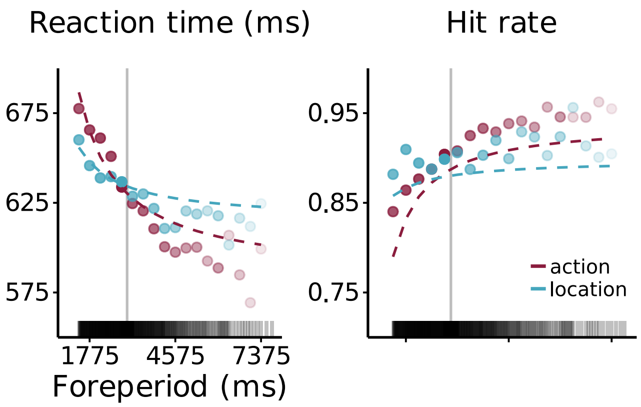


*Note.* Mean reaction time and hit rate as a function of action and location FP. Plotted as in Figure 5b.

### S.3 *f*MTP

All implementation details of the model can be found in the Python code (https://osf.io/eu7sd/, accompanied by a step-by-step tutorial). Salet et al. (2022) offers a detailed mathematical description of *f*MTP. Here, we only provide a qualitative description of the models as applied to WAM.

#### S.3.1 Specific versus Non-specific Preparation

In the main text, we characterized preparation in WAM as a specific process, meaning that each target presentation triggers its associated timing circuit that projects to a corresponding motor circuit representing a specific response (Figure 6b, main text). This model resembled the key aspects of the data (Figure 7). However, earlier work with *f*MTP has made the assumption that preparation reflects a general process independent of different response options (Salet et al., 2022). Here we present a version of the model that incorporates this assumption, and show that it fails to capture preparation effects observed in WAM (Figure S3a).

In this implementation, as for the specific implementation, each target onset still triggers one of three independent, timing circuits. However, the 'non-specific' motor circuit reflects inhibition and activation of *any* response. In both versions of the model, simultaneous activity in the timing and motor circuits drives Hebbian learning. In the 'specific' implementation of the main text, such learning only occurred between the two corresponding timing and motor circuits. The critical difference is that, since the 'non-specific' motor circuit represents preparation for any response option, such learning happens simultaneously for all three timing circuits.

#### S.3.2 Simulation

At each target presentation, the three timing circuits affect activity in the motor circuit. The amount of preparation for this target is determined as the summed preparation brought forth by the three timing circuits and directly translated to RT as explained in the main text. Figure S3 a shows the model's predicted RT as a function of

**Figure S.3**

*fMTP: non-specific implementation.*

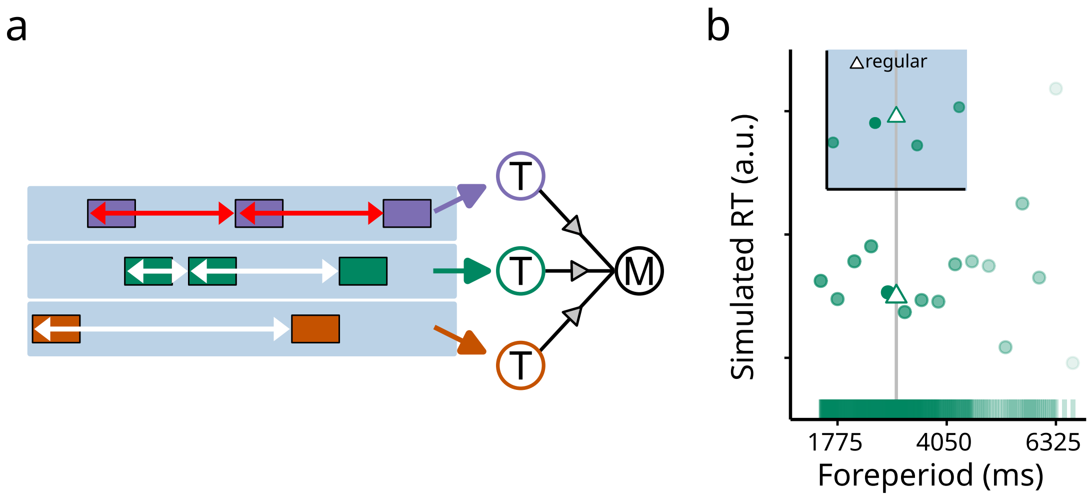


*Note.* (a) *f*MTP characterizing preparation as a non-specific process. Each of the timing circuits projects to the same motor circuit. (b) Mean simulated RT (arbitrary units, a.u.) as a function of FP (plotted as in Figure 7).

FP. Clearly, for this model, there is no indication that RT decreases as a function of FP as observed in the data.
